# Topographic Guidance in Melt-Electrowritten Tubular Scaffolds Enhances Engineered Kidney Tubule Performance

**DOI:** 10.1101/2020.09.16.300004

**Authors:** Anne Metje van Genderen, Katja Jansen, Marleen Kristen, Joost van Duijn, Yang Li, Carl C.L. Schuurmans, Jos Malda, Tina Vermonden, Jitske Jansen, Rosalinde Masereeuw, Miguel Castilho

## Abstract

To advance the engineering of kidney tubules for future implantation, constructs should be both self-supportive and yet small-sized and highly porous. Here, we hypothesize that the fabrication of small-sized porous tubular scaffolds with a highly organized fibrous microstructure by means of melt-electrowriting (MEW) allows the development of self-supported kidney proximal tubules with enhanced properties. A custom-built MEW device was used to fabricate tubular fibrous scaffolds with small diameter sizes (Ø = 0.5, 1, 3 mm) and well-defined, porous microarchitectures (rhombus, square, and random). Human umbilical vein endothelial cells (HUVEC) and human conditionally immortalized proximal tubular epithelial cells (ciPTEC) were seeded into the scaffolds and tested for monolayer formation, integrity, and organization, as well as for extracellular matrix (ECM) production and renal transport functionality. Tubular scaffolds were successfully manufactured by fine control of MEW instrument parameters. A minimum inner diameter of 0.5 mm and pore sizes of 0.2 mm were achieved. CiPTEC formed tight monolayers in all scaffold microarchitectures tested, but well-defined rhombus-shaped pores outperformed and facilitated unidirectional cell orientation, increased collagen type IV deposition, and expression of the renal transporters and differentiation markers organic cation transporter 2 (OCT2) and P-glycoprotein (P-gp). To conclude, we present smaller diameter engineered kidney tubules with microgeometry-directed cell functionality. Due to the well-organized tubular fiber scaffold microstructure, the tubes are mechanically self-supported, and the self-produced ECM constitutes the only barrier between the inner and outer compartment, facilitating rapid and active solute transport.

## 1. Introduction

*In vivo*, cells are surrounded by an extracellular matrix (ECM), a three-dimensional network composed of collagen, glycoproteins, polysaccharides, and other macromolecules [1]. Underneath epithelial and endothelial cells, including renal proximal tubules and peritubular vasculature, the ECM is formed as a thin, highly specialized basement membrane (BM), which acts both as supporting scaffold for the cells, and as barrier between an inner and outer compartment [1]. For kidney tubule engineering, a big challenge is the creation of renal proximal tubules in close proximity to peritubular vasculature. To allow for rapid exchange of solutes between blood and urine, the epithelial and endothelial BM should be virtually the only barriers between both cell types. To date, tubular tissue engineering mostly relies on non-porous, large diameter tubular scaffolds (Ø > 2 mm) for sufficient self-support, or smaller tubular lumens (Ø < 1 mm) surrounded by hydrogels to account for physiological functions, including vectorial transport [2-6]. The latter has been addressed recently in valuable *in vitro* tools for drug screening and disease modeling, with proven apical-basal cell polarity, and active reabsorption and transepithelial secretion function. These models provided evidence for the fact that tubular 3D curvature and biomimetic ECM properties enhance kidney cell functionality [2, 3, 5-8]. However, for the engineering of potentially implantable and durable kidney tubules, scaffolds must meet at least three criteria: they must be (1) small-sized and highly porous to increase the surface area, (2) freely accessible from the basolateral and luminal sides for rapid solute exchange and removal, and (3) flexible and yet strong enough to withstand intracorporeal forces such as pressure, tear and wear. Ideally, scaffolds for tubular tissue should also be resorbable and initially provide biophysical instructions to promote cell growth, differentiation, and BM production.

Fibrous scaffold meshes, alone or as reinforcement of hydrogels, can greatly improve the mechanical properties of engineered tissue. At the same time, they can influence cellular functionality and ECM deposition through proper scaffold design. Fibrous scaffolds have shown to facilitate cell migration and functionality by providing physiologically relevant mechanical stimulation [9-11]. For instance, cell growth and growth directionality can be positively affected by microfibers when ordered along a preferential direction [10, 12].

Previously, we provided proof-of-concept for stand-alone Ø < 1 mm kidney tubule grafts fabricated by solution electrospinning (SES), in which human kidney cells were able to bridge considerable fiber-to-fiber distances without the need of embedding hydrogels, which ensured rapid solute uptake [11]. Although SES was successful on manufacturing porous tubular fiber scaffolds, this process was rather limited by poor patterning control and hence fabrication reproducibility. Here, we fabricated tubular scaffolds using a superior electrohydrodynamic fiber printing technique called melt electrowriting (MEW) [13, 14]. MEW is a solvent-free technique that involves extrusion of a molten polymer fiber onto a computer-controlled collector. Organized 3D tubular constructs are obtained by precise fiber deposition on a rotating cylinder and successive layer-by-layer stacking, which require fine control over instrument parameters, in particular mandrel rotation and translation velocity [14-16]. Using MEW, we report the successful fabrication of Ø < 1 mm proximal tubule units that meet all above-mentioned criteria (Figure 1). The first aim was to determine optimal fabrication parameters for Ø = 0.5 mm and 1 mm tubular scaffolds and to assess their mechanical characteristics. The second aim was to apply these scaffolds to renal epithelial and vascular tubule engineering using conditionally immortalized proximal tubule epithelial cells (ciPTEC) and human umbilical vein endothelial cells (HUVEC); however, only renal epithelial cells were able to form monolayers within the scaffold pores and were used for subsequent experiments. Positive effects of tube structure and biomimetic materials on renal function have already been proven in literature; here, we hypothesized that by engineering tubular scaffolds with defined pore microarchitecture (*i*.*e*., rhombus, square and random), we can provide adequate topographic cues to promote cell directionality, autologous BM production, and renal transport function.

**Figure 1.**
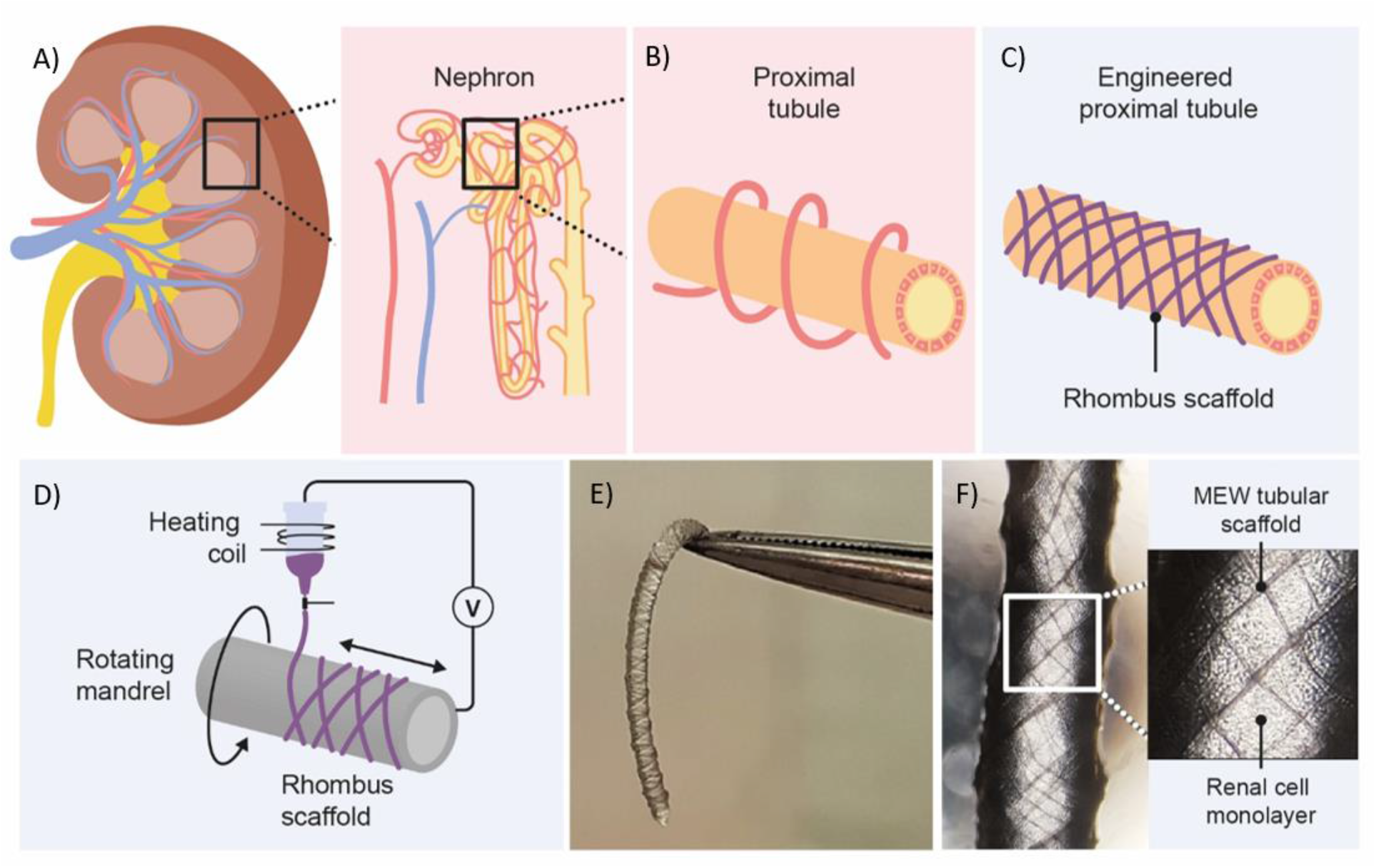
Graphical overview of design and fabrication of MEW medical grade polycaprolactone (PCL) tubular scaffolds. A) The kidney consists of roughly 1 million functional units, called nephrons. B) Active secretion takes mostly place in the proximal tubule, which is surrounded by peritubular capillaries. C-D) Melt-electrowriting was used to fabricate tubular scaffolds with defined pore geometries. E-F) Engineered kidney tubules contained confluent monolayers of proximal tubule epithelial cells.

## 2. Results

### 2.1. Design and Fabrication of Microfibrous Tubular Scaffolds

Fully resorbable medical grade polycaprolactone (PCL) tubular scaffolds were manufactured with a reproducible quality using an in-house built MEW set-up (Figure 1, 2, Movie S1). Scaffolds were fabricated with a range of winding angles, from 30° to 60°, by controlling the ratio between the tangential rotation velocity (V_tang_) and the translation velocity (V_trans_) (Figure S1). The measured winding angle was in accordance with the theoretical design (Figure 2A) and no significant differences in fiber diameter was observed when varying the winding angle (constant fiber diameter of approximately 20 µm, Figure 2B). Furthermore, while the winding angles could be accurately controlled for 3 mm and 1 mm mandrels, lower fidelity was observed for the 0.5 mm mandrel (Figure 2A-B). Additionally, the effect of fiber stacking on printed winding angle was evaluated (Figure 2C, Figure S2). In general, an increase in the deviation between designed and printed winding angle was observed when increasing the number of staked fibers from 1 to 10 layers. This difference was between 25% to 40% for 0.5 mm mandrels (Figure S2A) and below 10% for 3 mm (Figure S2B) and 1 mm mandrels (Figure 2C). Based on these printing fidelity results, tubes with 1 mm diameter and 10 layers (final thickness of approximately 100 µm) were selected for further experiments. Tubular scaffold with three different microarchitectures were fabricated, i.e. rhombus (winding angle 30°), square (winding angle 45°) and random. Notably, interconnected porosities above 90% and controlled pore sizes of 200 µm where obtained for rhombus (Figure 2D, 2E-F) and square pore shaped scaffolds, while only 40% porosity were observed for random scaffolds and no controlled pore size (Figure 2D).

**Figure 2.**
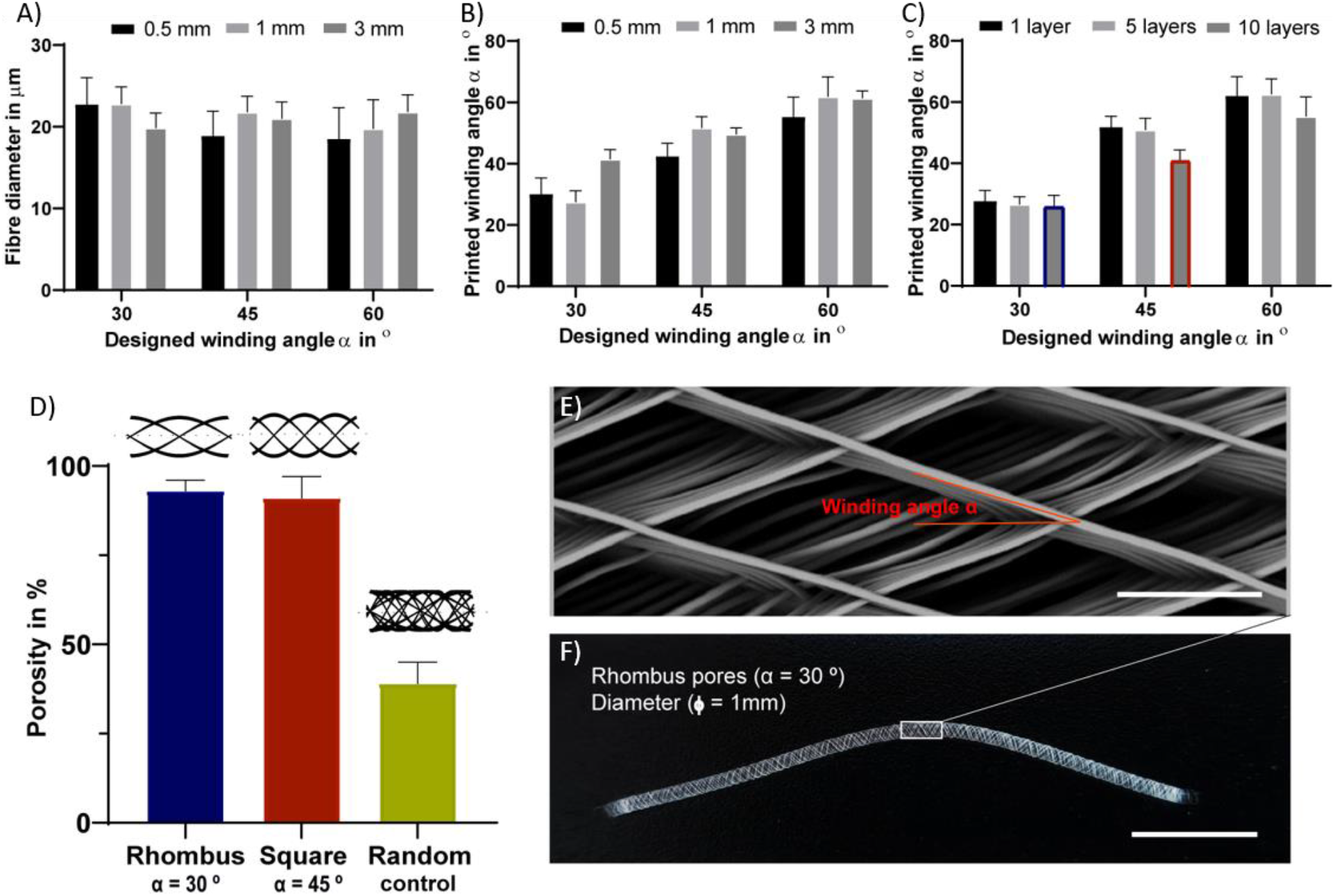
Tubular scaffold design and fabrication. A) variation of printed fiber diameters according to designed winding angles. B) Comparison between designed and printed winding angle used to assess printing accuracy. A-B were investigated for constructs with a single layer and for mandrels with three different diameters, 3, 1- and 0.5-mm. C) Comparison between designed and printed winding angle for constructs with 1, 5 and 10 stacked layers and mandrel with 1 mm in diameter. Blue and red identify rhombus and square constructs used for further mechanical and biological experiments. D) Porosity of fabricated tubes with 1 mm diameter. Representative E) stereoscopic and F) SEM image of printed tubular scaffolds with a rhombus (30°) microstructure for 1 mm mandrel. Scale bars = 100 µm (E) and 1 cm (F).

### 2.2. Cell Growth and Directionality

The first requirement of functional bioengineered kidney tubules is cell adherence and growth into a barrier-forming monolayer. Confirming our previous data, ciPTEC were able to form tight monolayers in L-DOPA biofunctionalized tubular scaffolds by stretching across the pores between the PCL microfibers [11]. L-DOPA coating increased scaffold hydrophilicity compared to non-coated tubular scaffolds (Table S1). Cells grew on the PCL-fibers, then they started bridging the pores from the corners and moved further inward; ultimately, the pores were entirely filled by ciPTEC monolayers. Only at a pore size of 400 µm and higher, ciPTEC were not able to bridge the gap.

Cell growth and filamentous actin (F-actin) orientation were studied in scaffolds with rhombus, square and random pore geometries (Figure 3A-I). For HUVEC, no microgeometry allowed monolayer formation; the cells only surrounded the PCL fibers (Figure 3A-C). When stained for platelet endothelial cell adhesion molecule 1 (PECAM-1 or CD31), a marker for endothelial cell junctional integrity, we observed irregular expression (Figure 3A-C). PECAM-1 is only expressed at the cell borders of confluent monolayers, whereas little or no expression is observed in sparse or migrating endothelial cells [17, 18]. In contrast to HUVEC, ciPTEC were able to fill the pores with all geometries investigated. For ciPTEC, rhombus-shaped pores caused preferential cell alignment along the scaffold fiber direction. Interestingly, the angle of deviation for ciPTEC grown in rhombus-shaped pores was approximately 25° as opposed to the 30° winding angle of the pores, indicating that ciPTEC are even more aligned than instructed by the geometry. The cell alignment order parameter *S* was calculated for ciPTEC, where a value of *S* = 1 would indicate that the long axis of the cells perfectly aligned with the scaffold fiber direction and *S* = 0 indicated no particular cell alignment. CiPTEC grown on rhombus-shaped pores exhibited cell alignment with an *S* of 0.51 ± 0.06, whereas both square and random microgeometries resulted in an isotropic distribution of cells with order parameters of 0 (Figure 3D-I) [19, 20].

**Figure 3.**
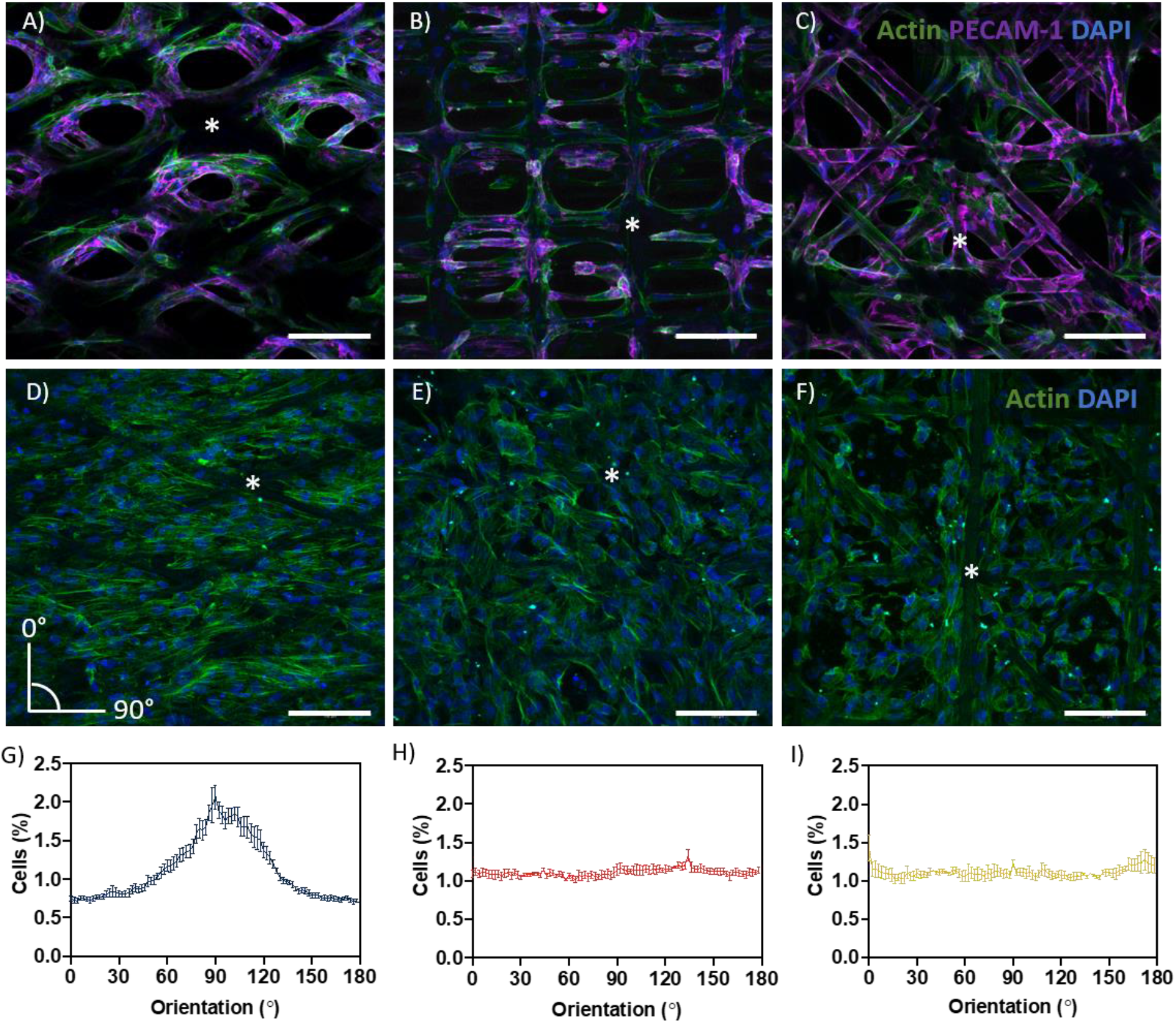
Cell growth and directionality. A-C) Actin filament (green), PECAM-1 (purple), and DAPI (blue) in A) rhombus-, B) square-, and C) random-shaped scaffolds for HUVEC. D-F) Actin filament (green) and DAPI (blue) in D) rhombus-, E) square- and F) random-shaped scaffolds for ciPTEC. Asterisks identify fiber scaffold position. G-I) Quantification of cell directionality in G) rhombus-, H) square- and I) random-shaped scaffolds for ciPTEC. Scale bars = 250 µm. Mean ± SEM, n=3.

### 2.3. Renal Transporter Expression and Function

The proximal tubule has a major role in waste secretion from the blood into the urine, and, therefore, expresses a broad range of functional transporters to facilitate this transport. We tested the effect of pore microgeometries on the gene expression of four relevant transporters: the efflux pumps breast cancer resistance protein (BCRP), multidrug resistance-associated protein 4 (MRP4), and P-glycoprotein (P-g), and the uptake transporter organic cation transporter 2 (OCT2). For all microgeometries, mRNA levels were compared to 2D culture (Figure 4A-D). For BCRP (Figure 4A, CT value 2D: 33.47 ± 0.12) and MRP4 (Figure 4B, CT value 2D: 29.60 ± 0.03), no differences were found. Remarkably, mRNA expression for P-gp increased 2.4-fold (Figure 4C, *p* < 0.05, CT value 2D: 29.43 ± 0.03) for tubular scaffolds with rhombus-shaped pores compared to 2D culture. Also, OCT2 expression was 3.4-fold higher (Figure 4D, *p* < 0.01, CT value 2D: 37.85 ± 0.30). In tubular scaffolds with isotropic pore microgeometries, transporter expression showed no significant increase compared to traditional 2D culture. Two functional assays were performed to validate increased transporter expression on functional level (Figure 4E-J). Calcein-AM diffuses into the cell, where it is hydrolyzed by esterases into fluorescent calcein, which is then secreted by BCRP and MRP4 (Figure 4E) [21]. When incubating the engineered kidney tubules with calcein-AM in presence of BCRP and MRP4 inhibitors KO143 and MK571, calcein accumulated in the cells (Figure 4G,I) (*p* <0.05). Rhodamine-123 (RH123) is a chemical fluorescent compound that is actively transported into the cell via OCT2 and actively excreted into the urinary compartment by P-gp and BCRP (Figure 4F). When the OCT2 inhibitor TPA was added, RH123 intracellular fluorescence intensity tended to be lower, although not significantly. When the P-gp inhibitor PSC-833 was added, intracellular fluorescence intensity increased (*p* < 0.05) (Figure 4I, J). Unfortunately, microscopic analysis was insufficiently sensitive to compare fluorescent intensities between engineered kidney tubules with different microgeometries (data not shown).

**Figure 4.**
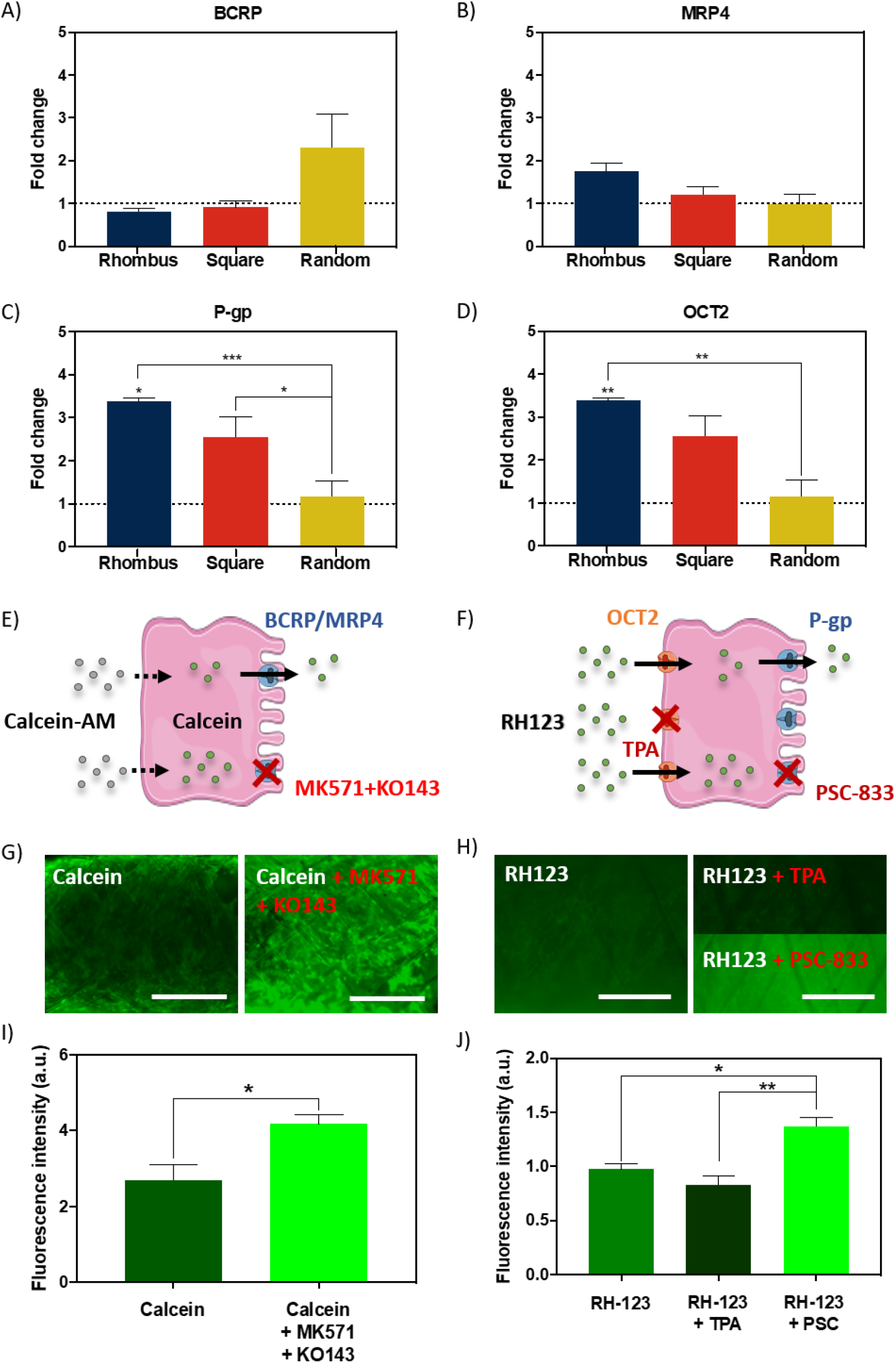
Renal transporter expression and function. A-D) Relative mRNA gene expression of ciPTEC cultured in tubular scaffolds compared to ciPTEC cultured in 2D (dotted line). E-F) Graphical overview of E) calcein-AM assay and F) rhodamine123 assay. G-H) Immunofluorescent images of G) calcein-AM assay and H) rhodamine123 assay in presence or absence of inhibitors. I-J) Quantification of immunofluorescent images measured as fluorescence intensity for I) calcein and J) rhodamine123. Scale bars = 500 µm. Mean ± SEM, n=3.

### 2.4. Basement Membrane Production

Collagen type IV is the most abundant component of the renal BM, making up 50% or more of the total [1]. To investigate how pore microgeometries affected ECM deposition, collagen type IV expression was measured over time, *i*.*e*., at day 7, 14, and 28, after starting maturation at 37°C (Figure 5A-B). In all pore microgeometries tested, ciPTEC deposited collagen type IV and filled all pores at the earliest timepoint measured (day 7). While collagen type IV deposition remained equal over time in random and square microgeometries, it significantly increased (*p* < 0.05) in rhombus scaffolds, with a maximum collagen type IV expression on day 28 (Figure 5B). On mRNA level, collagen type IV expression increased 5.5-fold in rhombus-shaped pores compared to traditional 2D culture at day 7 (*p* < 0.001, CT value 2D: 32.46 +/-0.33) (Figure 5C). Square-shaped pores also increased collagen IV mRNA levels in ciPTEC (4.4-fold, *p* < 0.01), whereas random-shaped pores did not have any beneficial effect compared to 2D cell culture (Figure 5C). Finally, we assessed how cells and their BM affected the mechanical properties of the MEW tubular scaffolds. Engineered kidney tubules were matured for 14 days before they were evaluated under uniaxial tensile loading to determine tangent modulus, yield strain, and elastic strain energy (Figure 5D-F, S3). A non-linear stress-strain behavior was observed for the different tested microarchitectures (Figure S3). Interestingly, without cells scaffolds with a rhombus geometry were able to reach higher elastic deformations (2-fold higher, approximately 10% deformation) and lower tangent modulus (1.7-fold lower, 4MPa) than scaffolds with random microstructure (Figure 5D-E, S3). No significant differences were observed on tangent modulus and elastic strain between scaffolds with a squared and random microarchitecture. In tubular scaffolds with ciPTEC monolayers, we observed a trend for an increase in yield strain (*p* = 0.0515) and an increase in elastic strain energy (*p* < 0.0002) in tubules with rhombus-shaped pores compared to scaffolds without cells (2-to 3-fold higher). It should be noted that a promising elastic behavior (shape recovery without visual permeant deformation) was observed for rhombus scaffolds after repetitive manual stretching, not observed for square and random tubular scaffolds (data not shown).

**Figure 5.**
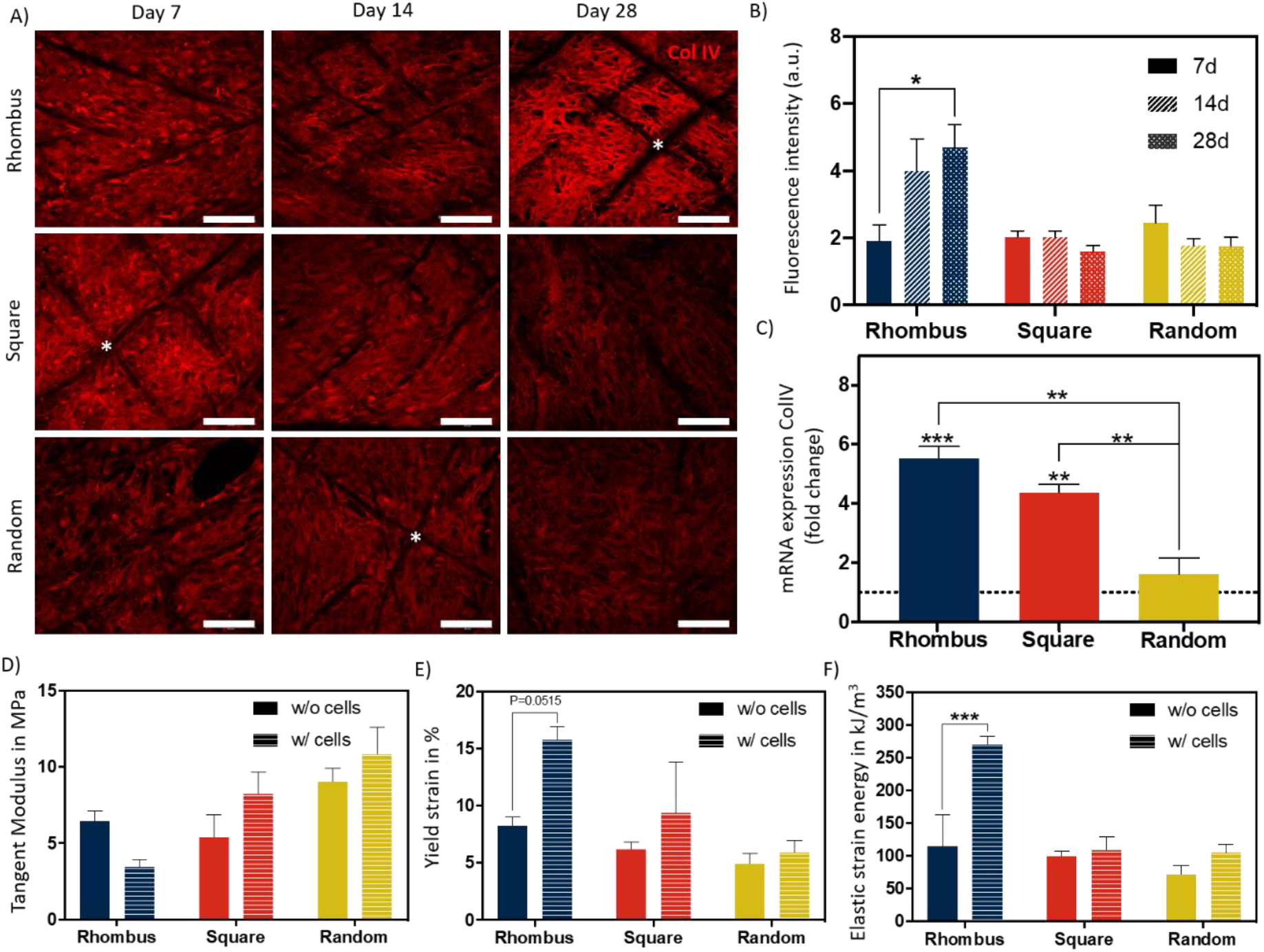
Basement membrane production and mechanical behavior of fibre constructs under tensile loading. A) Immunofluorescent images of collagen type IV (red) in rhombus-, square-, and random-shaped scaffolds captured at day 7, 14, and 28. Asterisks identify fiber scaffold position. Scale bars = 100 µm. B) Quantification of fluorescence intensity of the immunofluorescent images. C) Relative mRNA collagen type IV gene expression of ciPTEC cultured in tubular scaffolds compared to ciPTEC cultured in 2D (dotted line). D-F) Mechanical properties measured in tubular scaffolds with and without cells. Mean ± SEM, n=3.

### 2.5. Barrier Function

In our engineered kidney tubules, cells and their self-made BM deposited within the scaffold pores formed the only barrier between the basolateral and luminal side of the tubes. A thin but tight barrier is crucial for vectorial solute transport. To functionally assess barrier formation, inulin-FITC leakage was measured using a custom-made perfusion system. When mounting cell-free scaffolds in a dry chamber, inulin-FITC remained inside the scaffold due to surface tension (Figure 6A), but when PBS was added to the chamber, inulin-FITC directly leaked out (Figure 6B). In contrast, ciPTEC-laden scaffolds kept the inulin-FITC inside the lumen, indicating that the cells and their BM functioned as a leak-tight barrier between the luminal and basolateral sides of the engineered kidney tubule (Figure 6C). Engineered kidney tubules were also stained for collagen type IV and DAPI, and cross-sections were randomly made to confirm monolayer formation throughout the tubular construct (Figure 6D-E).

**Figure 6.**
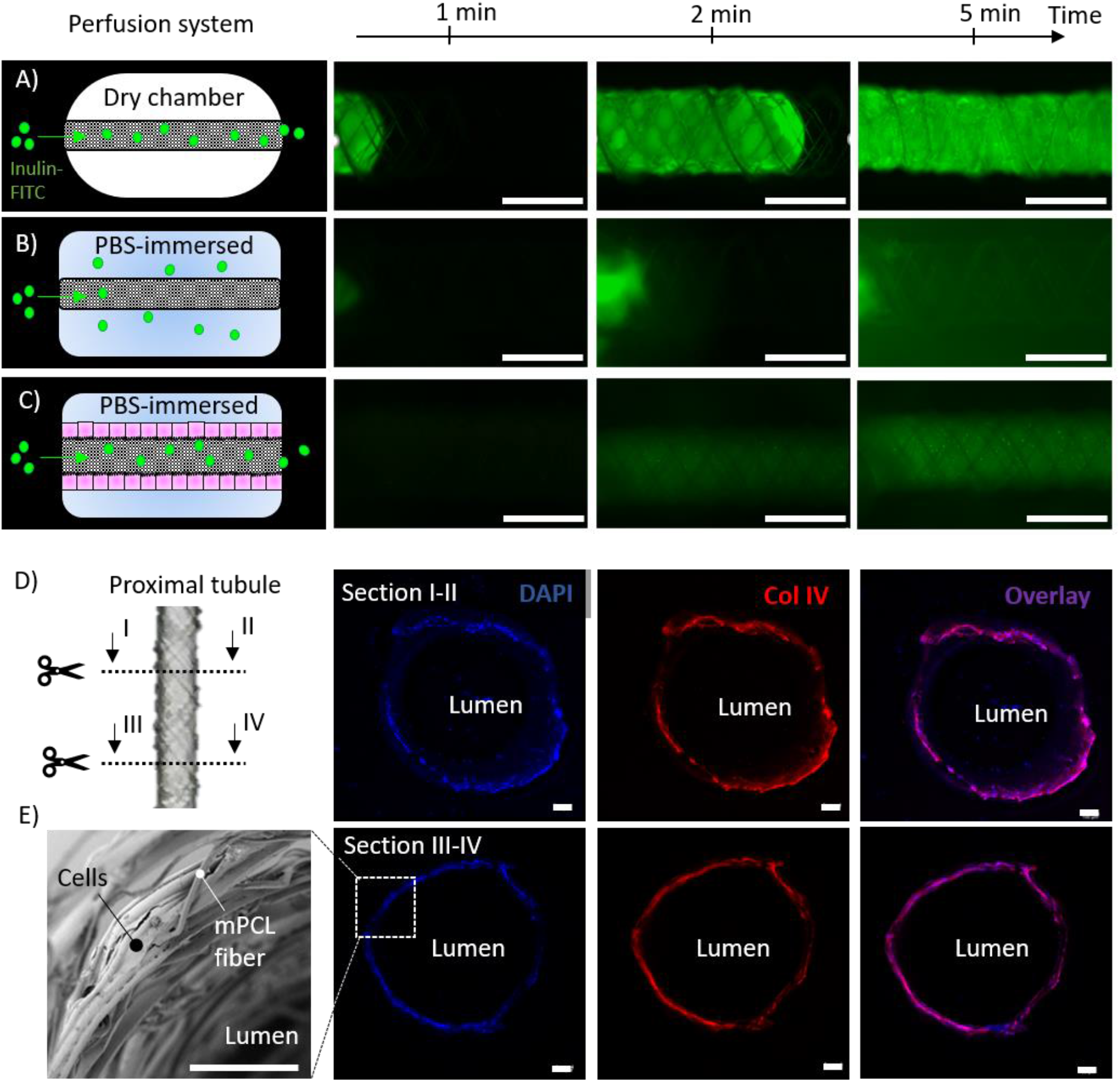
Renal barrier function. A-C) Inulin-FITC leakage assay in rhombus-shaped tubular scaffolds A-B) without and C) with cells. Perfusion chamber A) dry and B-C) immersed in PBS. Immunofluorescent images are captured after 1, 2 and 5 minutes of Inulin-FITC perfusion. D) Cross-sectional immunofluorescent images indicating a monolayer of ciPTEC throughout the construct. Immunofluorescent staining for DAPI (blue) and collagen type IV (red). E) Representative SEM image of cell laden rhombus-shaped tubular scaffolds demonstrating a renal monolayer. Scale bars = 100 µm.

## 3. Discussion

For rapid solute exchange and future implantation purposes, engineered kidney tubules must be flexible, wear-resistant, small in diameter, highly porous, and freely accessible from both the basolateral and luminal side. Using MEW, we fabricated self-supported, highly ordered tubular scaffolds with controlled pore microarchitectures (*i*.*e*., rhombus and square). We were able to produce scaffolds with an inner diameter of 0.5 mm, which is around 7 times the diameter of native kidney tubules and only 2 times the diameter of common dialysis fibers. While smaller diameters might be possible in future, the constructs also needed to remain easy to handle for experimental testing. The ability to control tubular scaffold microarchitecture is a unique characteristic of MEW that distinguishes the process from other conventional fiber formation technologies like SES. For kidney tubule engineering, we identified rhombus-shaped pores as a superior microgeometry over square-and random-shaped pores: topographic guidance resulted in directed cell growth, enhanced collagen type IV deposition, remarkably improved yield strain and elastic strain energy, and increased functional OCT2 and P-gp expression.

### 3.1. Rhombus-Shaped Pores Mimic *in vivo* Renal Tubular Basement Membrane Anisotropy Through Topographic Guidance

*In vivo*, the renal tubular BM exhibits an anisotropic structure, *i*.*e*., a directional preference in form of a polygonal meshwork of corrugated appearance [22]. In contrast to square-and random-shaped pores, rhombus-shaped pores guided cell growth along the scaffold fibers into a preferred cell direction with an angle of deviation that was even smaller than the winding angle of the scaffold. This phenomenon is called contact or topographic guidance [23, 24]. The effects of various topographic features on ciPTEC have been evaluated previously; already small features (<5 µm) could induce cell organization and alignment, which made the cells more susceptible to mechanical cues such as fluid shear stress (FSS) [25-27]. The alignment of the actin cytoskeleton might have contributed to the increased BM and transporter gene expression. Importantly, the cytoskeleton reorganization is known to alter force-activated pathways, as well as nuclear actin structures that can regulate gene expression and cell differentiation [28, 29]. Although more research is needed to confirm such underlying mechanisms in renal cells, our data do confirm beneficial effects of anisotropic cues on cell growth behavior and differentiation. The effects of different pore sizes and shapes of MEW scaffolds on cell alignment and gene expression have been shown to vary per cell type, including macrophages, bone marrow, smooth muscle cells, cardiomyocytes, and skeletal stem cells [30-34]. This indicates that the choice of pore size and shape need to be carefully considered depending on cell type of interest. For HUVEC, it has been shown that topographic cues of ridges and grooves can stabilize the endothelial phenotype. Based on the physiological BM microarchitecture *in vivo*, alignment in circumferential direction would be preferred [24, 35]. In our study, HUVEC failed in filling the MEW-fabricated pores of any microgeometry tested. Most likely, HUVEC have a smaller spreading length compared to ciPTEC, which is the only parameter besides initial pore size needed for predictions of void filling [36].

### 3.2. Renal Monolayers in MEW Pores Are Freely Accessible for Solute Uptake and Separate Compartments

Inhibitor-sensitive cellular uptake and excretion of fluorescent compounds demonstrated functional transport activity of our engineered kidney tubules. However, microscopic fluorescence uptake analysis was not sensitive enough to confirm increased expression of OCT2 and P-gp on a functional level. Vectorial solute transport requires an intact and polarized epithelial monolayer that forms a barrier between the luminal and the outer compartment. Microscopic images revealed consistent monolayer formation throughout the construct, which was confirmed by retainment of inulin-FITC during luminal perfusion. For uptake and subsequent drainage of metabolic waste products, an outside-oriented basolateral side, and an apical membrane facing the lumen are crucial. Usually, the interaction between cells and ECM are a determinant factor in the orientation and stabilization of cell polarity [37]. However, freshly seeded cells are not provided with a pre-existent ECM in our scaffolds, thus polarization likely depends on factors other than ECM cues. It is known that fluid shear stress (FSS) can facilitate cell polarization and has moreover been shown to enhance transporter expression in epithelial cells [2, 38-40]. Possibly, FSS could induce polarization and further enhance cellular functionality in our scaffolds, therefore future steps will include the establishment of a long-term flow-compatible cell culture system.

### 3.3. Rhombus-Shaped Pores Increase Yield and Elastic Strain Energy of Engineered Kidney Tubules

When handled or potentially implanted, engineered kidney tubules should be able to withstand external influences like pressure, tear and friction. When subjected to axial forces, controlled pore microgeometries themselves had a significant effect on tangent modulus, yield strain and elastic energy of tubular scaffolds when compared with non-controlled pore microgeometries. This evidence was even more pronounced when cells formed monolayers within the pores. Kidney tubules with rhombus-shaped pores could sustain higher strain (∼ 15%) before yielding compared to random and square (< 10%), probably due to the contact guided cell alignment with the fiber direction. Compared to tangent moduli of kidney tissue reported in literature (3-10 MPa), the tangent modulus of 4 MPa for the kidney tubules with rhombus-shaped pores (evaluated 14 days after monolayer formation) was close to the lower limit of this range [1]. However, considering the increase in ECM deposition over the course of 28 days, a higher tangent modulus can be expected over time. Notably, the control over scaffold microarchitecture on tubular constructs with 1mm allowed for aligned ECM deposition and consequent impressive compliance using PCL, a rather stiff thermoplastic material, as scaffolding material. It is important to mention, that future improvement of the MEW process is required to generate tubular scaffolds with tunable mechanical properties and internal diameters below 1 mm diameter. Recent works have reported the design and melt-electrowriting of auxetic tubular microfibers with unprecedent control of elastic deformations that could be applied in engineered kidney tubules. However, internal diameters obtained were one order of magnitude higher, 10mm, than the ones reported here [41].

### 3.4. Next Steps Towards Perfusable and Vascularized Engineered Kidney Tubules

As mentioned above, the establishment of long-term perfusion systems will further advance kidney tubule performance and readouts. Moreover, the flexible nature of MEW tubular scaffolds allows the intertwinement of multiple scaffolds, which would create direct contact between epithelium and endothelium, mimicking the physiological situation [1, 22]. Twisting scaffolds would additionally protect and stabilize the constructs. To engineer peritubular blood vessels, the initial pore size could be reduced by combination of MEW with electrospinning, as shown by Jungst *et al*. and Pennings *et al*. [42, 43]. Furthermore, the use of supportive cell types like pericytes or fibroblasts could help in the formation of endothelial monolayers [44]. In fact, direct contact between different cell types on PCL scaffolds, as opposed to indirect contact, can improve cell functionality [45]. Also, co-cultures of endothelial cells with renal epithelial cells have been shown to stimulate the balanced expression of various endothelial factors, which in turn improved the performance of renal cells [46].

In conclusion, we report the fabrication of self-supportive and yet small-sized porous MEW tubular scaffolds for kidney tubule engineering. MEW scaffolds enable direct access to the basolateral and luminal cell sides to facilitate solute exchange with vasculature in immediate proximity, which is critical for functional proximal tubule constructs. We showed that rhombus-shaped pores can enhance kidney tubule performance through topographic guidance, highlighting the fact that proper scaffold design offers opportunities to improve cell function.

## 4. Acknowledgements

A.M. van Genderen and K. Jansen contributed equally to this work. The Dutch Kidney Foundation (17PHD16) supported the work of A.M. van Genderen. The Netherlands Organization for Scientific Research (NWO) as part of the “Future Medicines” Program (022.006.003) supported the work of K. Jansen and C.C.L. Schuurmans. M. Castilho acknowledges the strategic alliance between University Medical Center Utrecht and Technical University Eindhoven. M. Castilho, J. Malda, T. Vermonden, and R. Masereeuw acknowledge the partners of Regenerative Medicine Crossing Borders (www.regmedxb.com), powered by Health∼Holland, Top Sector Life Sciences & Health.

## 5. Materials and Methods

### 5.1. Melt Electrowriting

We used a custom-built MEW device consisting of a rotating aluminum mandrel (Ø = 3, 1, 0.5 mm) mounted on a x-y axis and a custom print head mounted of a movable z-axis. The x-y-z axes were computer-controlled using an advanced motion controller MC403 (Trio Motion Technology Ltd.). Granular medical grade polycaprolactone (PCL, Purasorb PC 12, Corbion) was loaded into a 3 ml electrically heated glass syringe with a 27G size metallic nozzle and molten for 30 min prior printing. The molten polymer was electrified using a high voltage source (Heinzinger, LNC 30000-5 POS, 0–30 kV), and extrusion was assisted by a high-resolution air pressure regulator (VPPE-3-1-1/8-2-010-E1, Festo). In order to control scaffold microarchitectures, the effect of tangential rotation velocity (V_tang_) and translation velocity (V_trans_) on the laydown fiber winding angle (α) was investigated (Figure S1). The theoretical winding angle was determined according to,

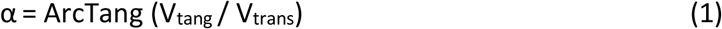

for 0 < α < 90°. The V_tang_ expressed in mm/s was assessed by,

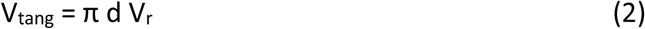

where d is the diameter of the collecting mandrel in mm and V_r_ is the rotational velocity of the mandrel in rad/s. Both V_tang_ and V_trans_ were varied between 0, 9, and 5.5 mm/s in order to obtain scaffolds with winding angles of 30 and 45°, corresponding to a square, rhombus, and random microarchitecture. The x-y position of the mandrel and respective translational velocity were programmed using a commercially available software package (Motion Perfect V4.2, Trio Motion Technology Ltd.) while the rotational velocity was controlled by Arduino IDE software. The high voltage, melting temperature, dispensing pressure and collecting distance between the printhead and mandrel were maintained in the ranges of 5-6.5 kV, 80-90] °C, 0.65-1 bar, and 4 mm, respectively for the fabrication of the three different scaffold microgeometries.

### 5.2. Imaging and Structural Characterization

Tubular scaffolds microstructure and fiber size were analyzed by stereomicroscope (Olympus SZ61) and by scanning electron microscopy (SEM, Phenom Pro, Phenom-World). SEM was performed at an acceleration speed of 5 -10 kV. Prior imaging, SEM scaffolds were gold sputtered coated (1nm) using a Q150R rotary-pumped sputter (Quorum Technologies). Images from both stereomicroscope and SEM were analyzed with Image J (National Instruments). Scaffold porosity was determined gravimetrically.

### 5.3. Cell Culture

Human conditionally immortalized proximal tubule epithelial cells (ciPTEC), isolated from human kidney tissue, were developed by Jansen *et al*. [47]. Briefly, proximal tubule cells obtained from healthy volunteers were infected with a temperature sensitive mutant U19tsA58 of SV40 large T antigen (SV40T) and the essential catalytic subunit of human telomerase (hTERT). ciPTEC were cultured at 33°C and 5% (v/v) CO_2_ up to 90% confluency to maintain a cell proliferation state. For maturation, ciPTEC were transferred to 37°C for at least 7 days prior to experimental readout. ciPTEC were cultured in T175 cultures flasks (Greiner Bio-One), using Dulbecco’s modified eagle medium/HAM’s F12 without phenol red (Thermofisher Scientific), supplemented with 5 µg mL−1 insulin, 5 µg mL−1 transferrin, 5 µg mL−1 selenium, 35 ng mL−1 hydrocortisone, 10 ng mL−1 epidermal growth factor, 40 pg mL−1 tri-iodothyronine (Sigma-Aldrich), 10% fetal bovine serum (FBS) (Greiner Bio-One), and 1% penicillin/streptomycin (Thermofisher Scientific). Primary human umbilical vein endothelial cells (HUVEC) (American Type Culture Collection (ATCC)) were cultured using vascular cell basal medium and endothelial cell growth kit-VEGF (ATCC). HUVEC were cultured in T175 culture flasks (Greiner Bio-One) at 37°C and 5% (v/v) CO_2_, and confluency was maintained at a maximum of 90% throughout the culture period.

### 5.4. Scaffold Coating and Cell Seeding onto Scaffolds

Before cell seeding, PCL tubular scaffolds were sterilized using 70% ethanol evaporation and UV exposure (365 nm for 15 min). 2 mg/mL L-3,4-dihydroxyphenylalanine (L-DOPA, Sigma-Aldrich) was dissolved in 10 mM tris(hydroxyethyl)aminomethane (Tris) pH 8.5 buffer at 37°C for 45 min [7, 48, 49]. After sterile filtration (0.2 µm pore size), scaffolds were coated through submersion for 4h at 37°C and thoroughly washed with phosphate buffered saline (PBS without calcium and magnesium, Lonza). Next, 10 × 10^6^ cells/ml were added in small culture medium drops onto flat scaffolds or injected into tubular scaffolds. Cells were incubated for 4h at 33°C (ciPTEC) or 37°C (HUVEC); tubular scaffolds were turned 180° after 2h. Afterwards, culture medium was added, and the constructs were cultured until confluency at 33°C and then at 37°C (ciPTEC), or directly at 37°C (HUVEC). Scaffolds were cultured for 1-4 weeks at 37°C. Culture medium was refreshed 3x per week. Changes on polymer surface wettability before and after L-DOPA coating were evaluated by static contact angle measurements using sessile drop technique (Data Physics, OCA 15EC). Measurements were performed according to a procedure described elsewhere [10].

### 5.5. Immunofluorescence

Scaffolds were fixed for 5 min with 4% (v/v) paraformaldehyde in PBS and permeabilized with 0.3% (v/v) triton X-100 in PBS for 10 min at RT. Subsequently, cells were exposed to blocking buffer consisting of 2% (v/v) FCS, 0.5% (w/v) bovine serum albumin (BSA), and 0.1% (v/v) Tween20 in PBS for 30 min at RT. Antibodies were diluted in blocking buffer. Primary antibodies were incubated O/N at 4°C and secondary antibodies for 1h at RT. We used the following primary and secondary antibodies and dilutions: mouse anti-α-tubulin 1:150 (Sigma-Aldrich), goat anti-collagen IV 1:50 (Southern Biotech), and mouse anti CD31 1:250 (Abcam), AlexaFluor 488 donkey-anti-rabbit 1:200 (Invitrogen, lot# 1981155), AlexaFluor 488 goat-anti-mouse 1:200 (Invitrogen), and AlexaFluor 647 donkey-anti-goat 1:200 (Invitrogen). Moreover, we used AlexaFluor 546 phalloidin 1:1,000 (Thermo Fisher Scientific) to stain actin filaments, and nuclei were stained using DAPI 1:1,000 (Sigma-Aldrich). Immunofluorescence was examined using confocal microscopy (Leica TCS SP8 X) and software Leica Application Suite X. Images were analyzed using ImageJ. Images were converted to 8-bit, Z-projections were made, and the same threshold was applied for every image. Actin filament directionality was quantified using the directionality functionality and Fourier components analysis.

### 5.6. Gene Expression

Gene expression analysis was performed by RT-qPCR on ciPTEC after growing at 33°C until confluency and 7 days at 37 °C. RNA was isolated from ciPTEC grown in tubular scaffolds or in L-Dopa coated well plates (control) using a PureLink RNA mini kit (Invitrogen), and cDNA was prepared using 500 ng sample RNA templates using iScript cDNA Synthesis Kit (BioRad). RT-qPCR analysis was performed on a CFX96 real-time PCR detection system (BioRad) using TaqMan Universal PCR Master Mix (Invitrogen) and TaqMan Gene Expression Assay for COL4A1, ABCG2, ABCC4, ABCB1 and SLC22A2 (Invitrogen). All reactions were carried out with equivalent dilutions of each cDNA sample.

### 5.7. Mechanical Testing

The mechanical properties of cell free and cell cultured tubular constructs (day 14 of culture at 37°C) were studied under uniaxial tensile test using a Dynamic Mechanical Analyzer (DMA Q800, TA Instruments). Before testing, cell cultured constructs were washed with HBSS, fixated for 10 minutes with 4% (v/v) paraformaldehyde and then washed again with HBSS. Tensile tests were performed at a rate of 0.1 N/min. Prior testing, nominal diameter and length of each construct was measured using stereomicroscopic and SEM images. The tangent modulus, yield strain and elastic strain energy were determined from engineered stress-strain curves. Tangent moduli were determined using a least square fit of the initial slope of the stress-strain elastic region (E_T_, as presented in Figure S3), yield point (ε_el_) was defined as point where nonlinear deformation begins and elastic strain energy (U_0_) was obtained,

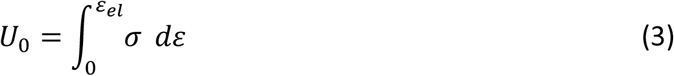

where σ and ε represent engineered stress and strain, respectively.

### 5.8. Functional assays

To test the functionality of various transporters (BCRP, MRP4, P-gp and OCT2), tubular scaffolds with ciPTEC were matured for 3 weeks, rinsed in HBSS, and incubated with fluorescent substrates. Cells were incubated with 1 μM non-fluorescent calcein-AM for 15 min at 37°C to confirm cellular metabolism through enzymatic conversion of calcein-AM to fluorescent calcein. Incubation was performed in presence or absence of 5 μM MK-571 and KO173 (inhibitors of MRP4 and BCRP) to validate functional excretion of calcein. Moreover, tubular scaffolds were incubated with 10 μM rhodamine123 (RH123), substrate for uptake transporter OCT2 and excretion transporter P-gp, in presence or absence of 5 μM tetrapentylammonium (TPA^+^, OCT2 inhibitor) or 5 μM PSC-833 (P-gp inhibitor) for 40 min at 37°C. After incubation, the scaffolds were rinsed with HBSS and images were captured with green filters using the Keyence BZ-9000 fluorescence microscope (Keyence). ImageJ (National Instruments) was used to measure fluorescence intensity in 16-bit images after background subtraction. In addition, a leakage assay was performed to assess engineered proximal tubule function. First, a customized perfusion chamber was designed and printed with polylactic acid (PLA) using a fused filament fabrication method (Ultimaker). The printed chamber was mounted on a glass slide using epoxy glue (Loctite), and tubular scaffolds were fixed inside with dental glue (GI-MASK Automix, Coltene) (Figure S4). Scaffolds were perfused with 0.1 mg/ml fluorescein isothiocyanate-inulin for 5 min (inulin-FITC, Sigma-Aldrich), and real-time imaged using a fluorescence microscope with a green filter (Keyence BZ-9000).

### 5.9. Statistical Analysis

Unless otherwise stated, 3 scaffolds per condition were used in 3 independent experiments respectively. Statistical analysis was carried out in Graphpad Prism 8.1.0 (GraphPad Software Inc.) using a student t-test when two groups were compared, and a one-way ANOVA and Tukey’s multiple comparisons post-hoc test when 3 or more groups were compared. A two-way ANOVA and multiple comparisons were used for comparison of multiple groups with 2 independent variables. Differences were considered significant with a p-value of p < 0.05. * indicates p < 0.05, ** p < 0.01, *** p < 0.001 and **** p < 0.0001. Data are represented as mean ± standard error of the mean (SEM) or as mean ± standard deviation (SD).

## Supporting Information

**Table S1:**
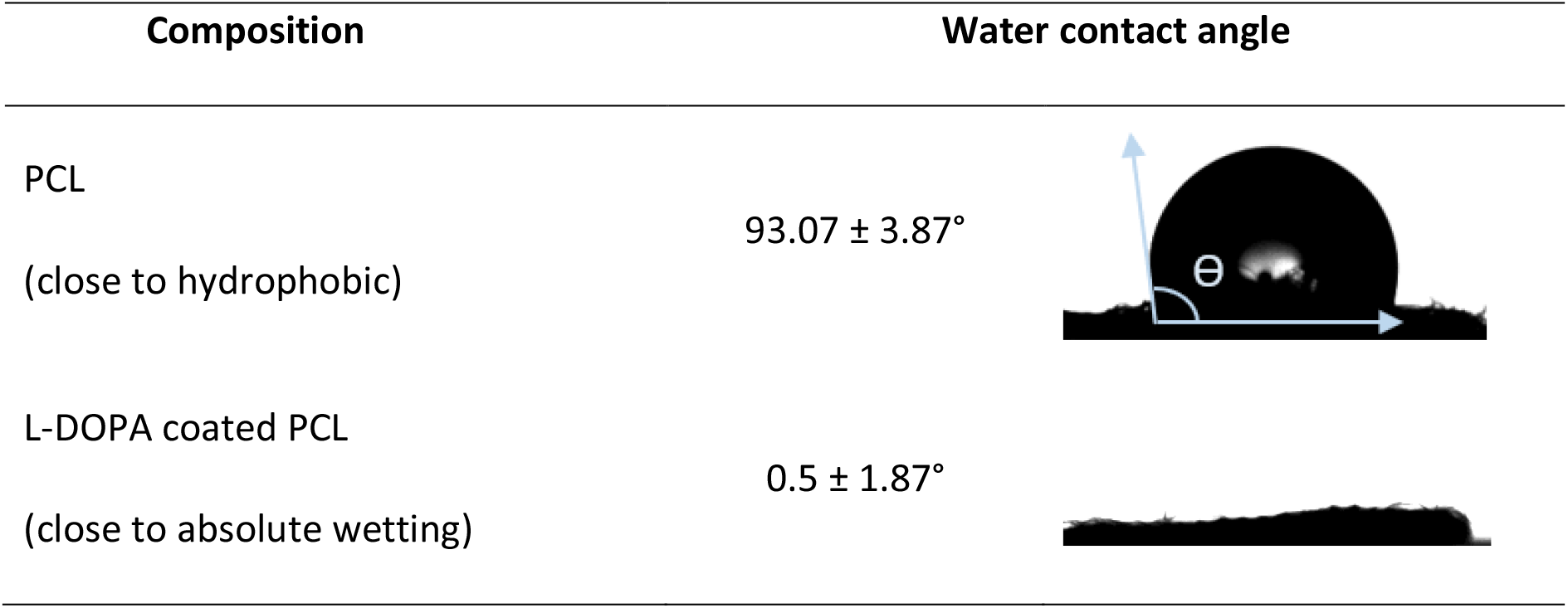
Water contact angles on uncoated PCL and L-DOPA coated PCL surfaces. Mean ± SEM, n=3.

| Composition | Water contact angle |  |
| --- | --- | --- |
| PCL<br>(close to hydrophobic) | $93.07 \pm 3.87^\circ$ | |
| L-DOPA coated PCL<br>(close to absolute wetting) | $0.5 \pm 1.87^\circ$ | |

**Figure S1:**
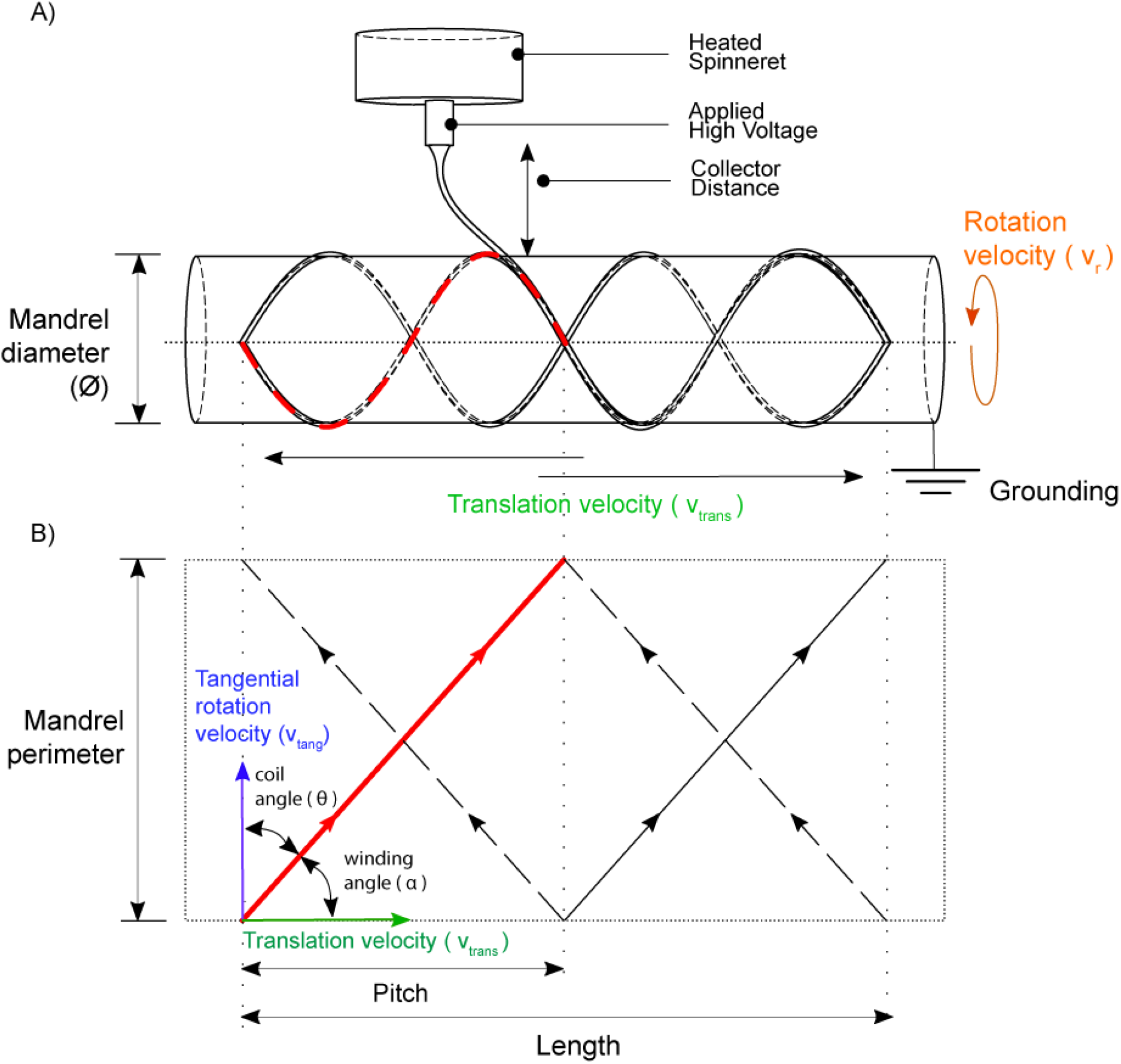
A) Schematic of the melt electrowriting set-up, showing the collection of thin electrified polymer jets onto a computer controlled, rotating and translating mandrel. B) Representation of the mandrel velocities, translational (V_trans_), tangential (V_tang_), rotational (V_rot_), as well as winding angle (α), coil angle (⊖), tube length, pitch and mandrel perimeter (r) in the mandrel surface during fiber collection. Winding angle was engineered by the ratio between V_tang_ and V_trans_ at constant fiber length.

**Figure S2:**
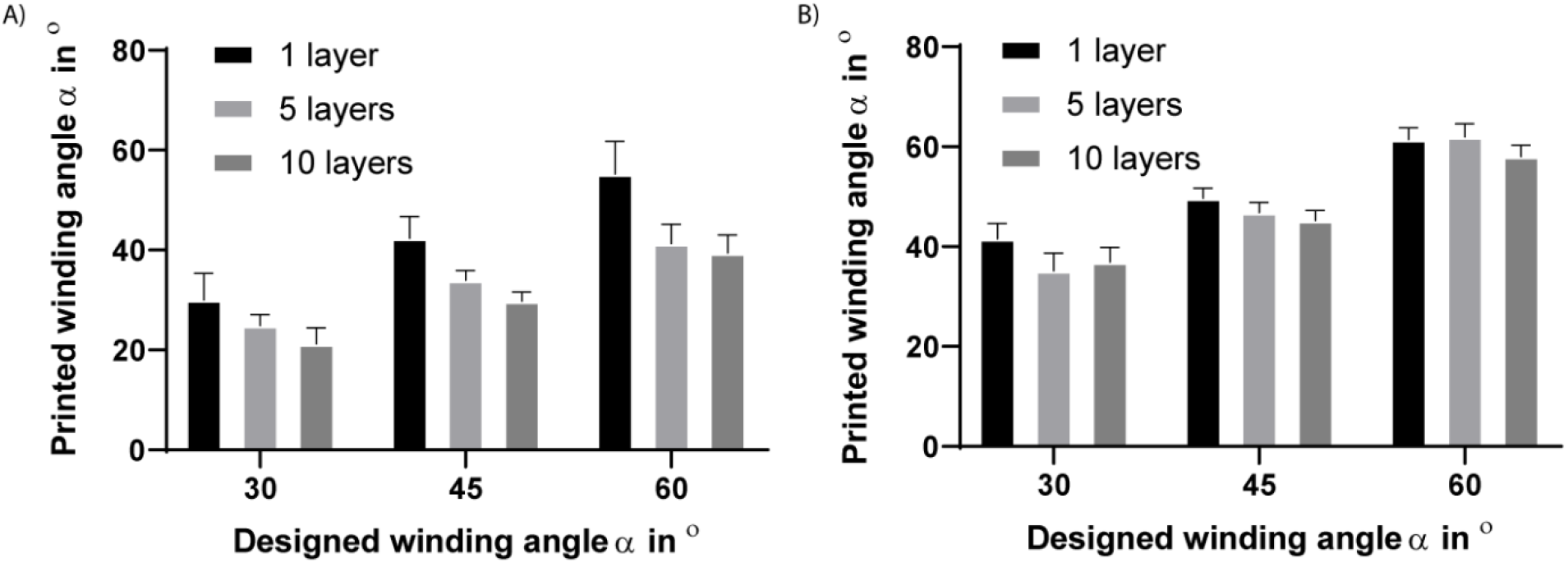
Comparison between designed and printed winding angle for constructs with 1, 5 and 10 stacked layers and mandrel with A) 0.5 mm and B) 3mm in diameter.

**Figure S3:**
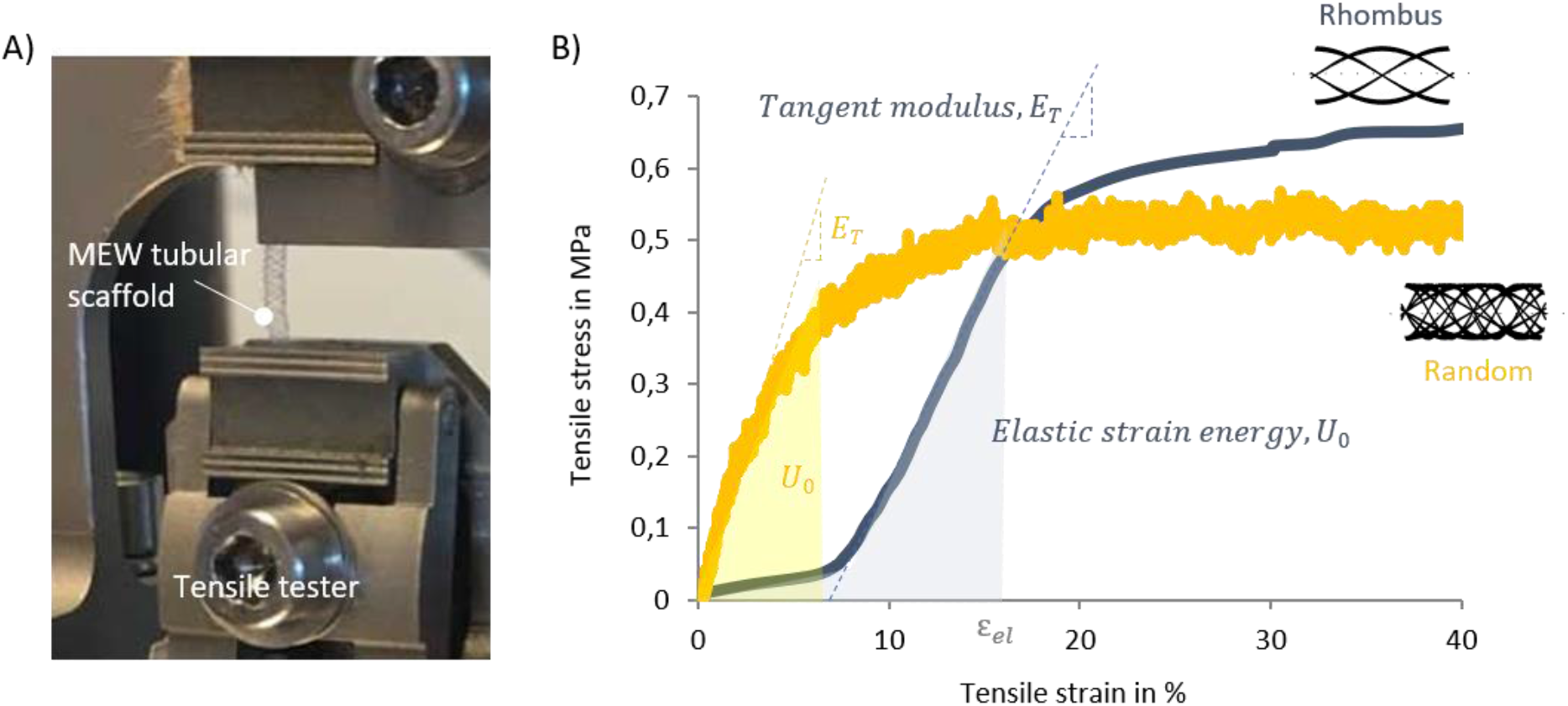
A) Tensile tester set up and B) representative engineered stress-strain curves with determined mechanical parameters for tubular scaffolds with rhombus and random microarchitecture with cells.

**Figure S4:**
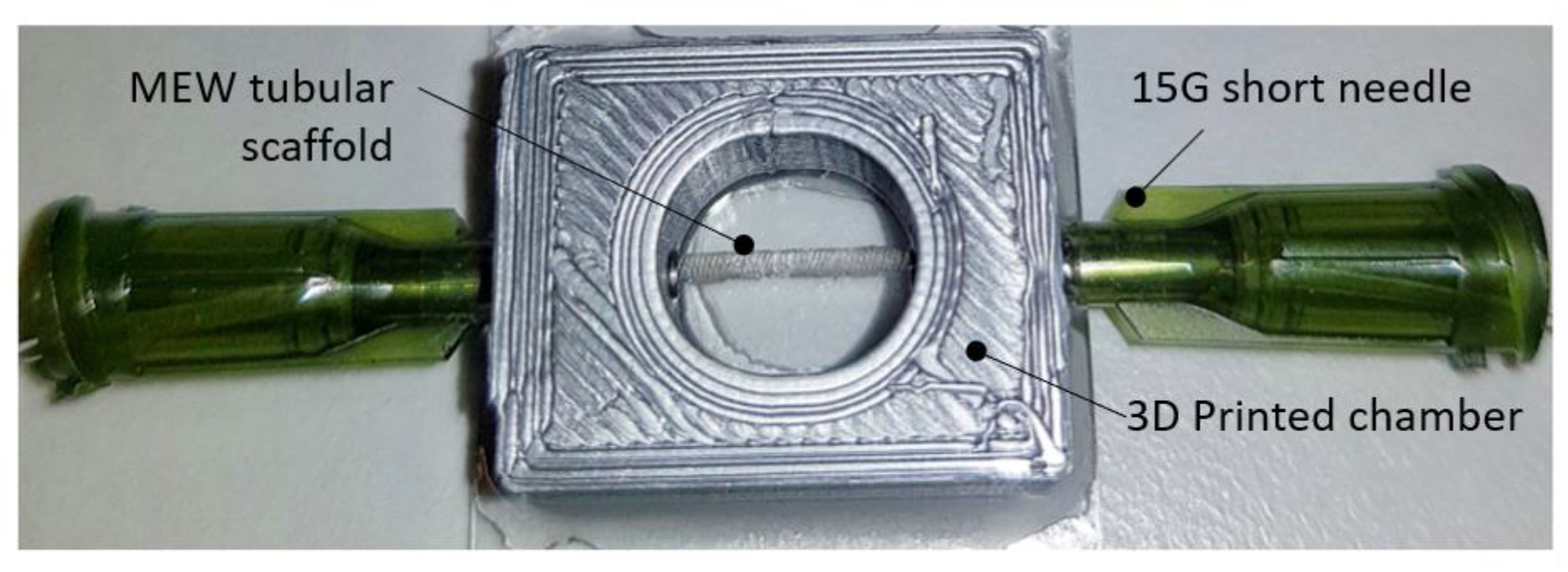
Custom made perfusion system used for renal barrier function studies. MEW tubular scaffolds were mounted in a 3D printed PLA customized chamber and then perfused.

## Movies

**Movie S1**. Video for melt-electrowriting of tubular constructs with rhombus microstructure

